# Prefrontal cortical protease TACE/ADAM17 is involved in neuroinflammation and stress-related eating alterations

**DOI:** 10.1101/2023.01.23.525269

**Authors:** Fransua Sharafeddin, Mina Ghaly, Timothy B. Simon, Perla Ontiveros-Ángel, Johnny D. Figueroa

## Abstract

Childhood traumatic stress profoundly affects prefrontal cortical networks regulating top-down control of eating and body weight. However, the neurobiological mechanisms contributing to trauma-induced aberrant eating behaviors remain largely unknown. Traumatic stress influences brain immune responses, which may, in turn, disrupt prefrontal cortical networks and behaviors. The tumor necrosis factor alpha-converting enzyme / a disintegrin and metalloproteinase 17 (TACE/ADAM17) is a sheddase with essential functions in brain maturation, behavior, and neuroinflammation. This study aimed to determine the role of TACE/ADAM17 on traumatic stress-induced disruption of eating patterns. We demonstrate a novel mechanistic connection between prefrontal cortical TACE/ADAM17 and trauma-induced eating behaviors. Fifty-two (52) adolescent Lewis rats (postnatal day, PND, 15) were injected intracerebrally either with a novel Accell™ SMARTpool ADAM17 siRNA or a corresponding siRNA vehicle. The RNAscope Multiplex Fluorescent v2 Assay was used to visualize mRNA expression. Observation cages were used to monitor ethological behaviors in a more naturalistic environment over long periods. We found that traumatic stress blunts startle reactivity and alter eating behaviors (increased intake and disrupted eating patterns). We also found that the rats that received prefrontal cortical TACE/ADAM17 siRNA administration exhibited decreased eating and increased grooming behaviors compared to controls. These changes were associated with decreased AIF-1 expression (a typical marker of microglia and neuroinflammation). This study demonstrates that prefrontal cortical TACE/ADAM17 is involved in neuroinflammation and may play essential roles in regulating feeding patterns under stress conditions. TACE/ADAM17 represents a promising target to ameliorate inflammation-induced brain and behavior alterations.

## INTRODUCTION

In the United States, 60% of children have been exposed to at least one traumatic event.^1^ Around 40% of high school students have experienced violence, and up to 6% are diagnosed with post-traumatic stress disorder (PTSD).^2,3^ As such, pediatric PTSD is an emerging and significant public health problem. PTSD has been associated with a higher body mass index (BMI) and obesity and its consequential metabolic complications.^4,5^ Changes in the hypothalamic-pituitary-adrenal (HPA) axis homeostasis may account for increased food intake and obesity in individuals exposed to stress.^6^ However, recent evidence suggests that brain regions responsible for the cognitive control of food intake may override homeostatic processes to promote food intake and weight gain.^7^

The prefrontal cortex (PFC) is a cortical region with the most substantial growth during development. It comprises almost one-third of the adult human neocortex.^8^ Undergoing significant expansion during maturation, the PFC shows a more extended course of formation than other cortical regions.^9^ The PFC has different areas, including the dorsolateral, orbitofrontal, and ventromedial area.^10^ The PFC is central in mediating cognition and behavior.^11^ Specifically, this brain region is associated with the higher-order cognitive and social-emotional functions and is responsible for conducting complex goal-directed activities representing the executive function.^12^ In particular, the medial prefrontal cortex (mPFC) is implicated in cognitive function, social and feeding behaviors, food valuation, motivation, and emotional regulation.^13^ The capability for complete functional control is supported by extensive neuronal networks connecting the PFC to different regions.^14^ Due to its extended course of maturation compared to other cortical regions, the mPFC is exceedingly vulnerable to traumatic stress exposures during childhood. Traumatic stress leads to abnormal development of the mPFC, which has been associated with eating disorders and obesity.^15^ Despite recognizing childhood trauma as a significant risk factor for eating disorders and obesity, it is poorly understood by which molecular mechanisms trauma exposure during childhood alters mPFC maturation and function.^16-18^

The formation of synaptic networks continues postnatally and represents a complex synaptogenesis and synaptic pruning process that occur concurrently and shapes mPFC circuits.^19^ Synaptic refinement involves microglia, which are the innate immune cells of the brain. Microglia play a critical role in shaping the PFC, particularly during adolescence.^20,21^ This process is tightly regulated through intricate permissive and repulsive growth pathways that drive molecular signaling and synaptic maturation. The tumor necrosis factor alpha-converting enzyme / a disintegrin and metalloproteinase 17 (TACE/ADAM17) is one of the main sheddases required for the cleavage of different growth factors and inflammatory mediators.^22^ TACE/ADAM17 has been implicated in various diseases, including heart failure, diabetes, cancer, atherosclerosis, arthritis, and central nervous system pathologies.^23,24^ Because of its essential role in microglial survival and phagocytic functions,^25^ we recently proposed a model in which supraoptimal TACE/ADAM17 activities may contribute to neuroinflammation and altered brain maturation.^26^ Since neuroinflammation plays a critical role in mPFC maturation and function, we reasoned that TACE/ADAM17 might contribute to behavioral alterations associated with stress-induced eating and obesity.^27-29^

## MATERIALS AND METHODS

### Rat Model

All the experiments were performed following animal protocol 20-171, approved by the Institutional Animal Care and Use Committee (IACUC) at the Loma Linda University School of Medicine. Animals were kept in typical housing conditions (21 ± 2 °C, relative humidity of 45%, 12-hour light/dark phases with lights on at 7:00 AM, paired housed for control groups). The body weights were recorded weekly or daily during the week of behavioral testing. Food consumption was quantified at least twice per week. The rats were never food or water restricted. In this study, we used Lewis rats, which have been shown to possess decreased activation of the hypothalamic-pituitary-adrenocortical (HPA) axis in response to stress. These characteristics mimic neurophysiological processes in humans exposed to trauma, making this rat model suitable for our proposed experiments.^30,31^ We and others have used Lewis rats to model abnormal neuronal maturation in adolescence due to obesogenic diet exposure;^32^ genetic influence on addiction vulnerability, with particular emphasis on differences in mesolimbic dopamine transmission, rewarding and emotional function;^33^ high-fat diet-induced increase in susceptibility to traumatic stress during adolescence.^34^

### Traumatic Stress Model

The traumatic psychosocial stress (PSS) protocol was adapted from an established rat model of traumatic stress. PSS produces cognitive impairments and maladaptive behaviors lasting up to 4 months.^35,36^ The PSS consisted of two exposures to a cat that lasted one hour each while the animal was immobilized. Exposures took place on days 1 and 10 of the PSS. The animals in the stress group underwent social isolation, composed of single housing during the stress perturbation and experimentation period. This model mimics critical constituents of trauma in humans, including lacking control over stressful situations and the inability to predict upcoming events. It also assimilates loneliness, social isolation, and lack of social support, essential psychosocial components of PTSD.

### Experimental Groups

Fifty-two (52) adolescent Lewis rats (postnatal day, PND, 15) were acquired from Charles River Laboratories (Portage, MI). Animals were habituated to housing conditions with 12-h light/dark phases for at least one week before the initiation of the experiments. The rats were weaned at PND 22. Rats were matched by sex, body weight, and startle reactivity. Subsequently, the rats were assigned into **six** groups based on trauma exposure and treatment: **1)** Naïve + Unexposed (*n* = 8), **2)** Naïve + PSS (*n* = 8), **3)** Control + Unexposed (*n* = 12), **4)** Control + PSS (*n* = 8); **5)** siRNA + Unexposed (*n* = 10), **6)** siRNA + PSS (*n* = 13). Unexposed and naïve groups were housed in pairs (same-sex and treatment). Both sexes were included in each group (half males and half females). The rats were given *ad libitum* access to water and a purified diet (product no. F7463; Bio-Serv, Frenchtown, NJ). This diet was used to provide continuity to our studies. Food consumption was monitored, and body weight was measured every week. While classic behavioral readouts were acquired during the light phase, naturalistic behaviors were examined during long periods (24-48 h), encompassing light and dark phases. The study timeline is summarized in **Figure 1**.

**Figure 1.**
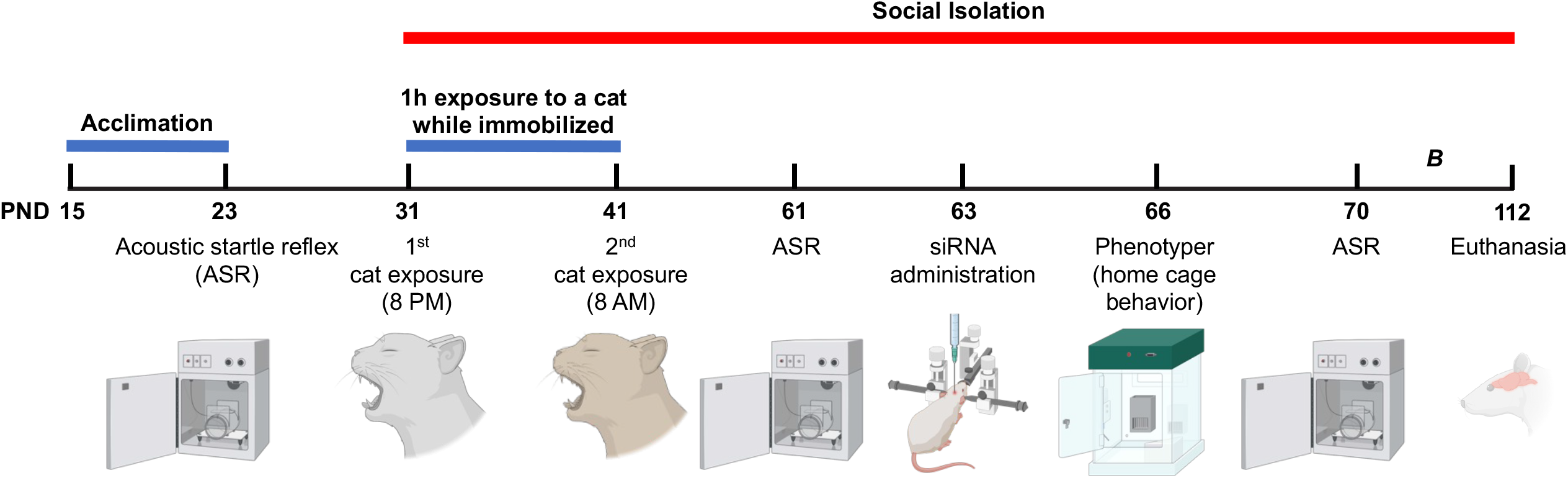
Study design and timeline of experimental procedures, behavioral tests, and outcome measures. Adolescent rats were matched based on their acoustic startle reflex (ASR) responses and allocated to one of the six groups: naïve unexposed (Naïve UNEX), vehicle control unexposed (Control UNEX), siRNA unexposed (siRNA UNEX), naïve exposed (Naïve EXP), vehicle control exposed (Control EXP), and siRNA exposed (siRNA EXP). The traumatic psychosocial stress (PSS) protocol consisted of two exposures to a cat that lasted one hour each while the animals were immobilized. Exposures were on days 1 and 10 of the PSS. The animals in the exposure group underwent social isolation composed of single housing during the experimentation period. The ASR experiments were performed before the beginning of the PSS protocol (PND23), before (PND61) and after (PND70) siRNA surgeries, and before euthanasia (PND107). The siRNA injection was performed on PND63. For behavioral assessments, we evaluated home cage behaviors in the Phenotyper on PND66. Additional long-term outcomes were examined, including high-fat diet food intake and social behaviors (B; see Figure 5). All the rats were euthanized on PND112.

### RNA Interference

A novel Accell SMARTpool ADAM17 siRNA was used (Cat# E-080034-00-0050, Horizon Discovery, Lafayette, CO, USA), which requires no transfection reagent or viral vector for delivery. Four different sequences targeting the ADAM17 gene were pooled: 1) Accell SMARTpool siRNA A-080034-13, Adam17, Target Sequence: GGAUUAGCUUACGUUGGUU, molecular weight: 13,563.8 (g/mol), extinction coefficient: 356,534 (L/mol·cm); 2) Accell SMARTpool siRNA A-080034-14, Adam17, Target Sequence: GUAUAAGUCUGAAGAUAUC, molecular weight: 13,495.7 (g/mol), extinction coefficient: 371,664 (L/mol·cm); 3) Accell SMARTpool siRNA A-080034-15, Adam17, Target Sequence: UCAUCGAUUUUAUAAGUAC, molecular weight: 13,485.8 (g/mol), extinction coefficient: 375,224 (L/mol·cm); 4) Accell SMARTpool siRNA A-080034-16, Adam17, Target Sequence: UUAUGGAGUACAGAUAGAA, molecular weight: 13,440.6 (g/mol), extinction coefficient: 370,062 (L/mol·cm).

### Surgery

Animals were transported to the surgery room 30 min before the surgical procedure. First, the rats were placed into an anesthesia induction chamber, and the Isoflurane dosage was set to 4 liters per minute (lpm). After anesthesia induction, the rats were shaved and placed into the stereotactic device, and ear pins were placed. A midline incision was made, and a bur hole was placed on the right side corresponding to the right mPFC. Stereotactic coordinates for needle insertion were 3.5 mm anterior, 0.6 mm lateral, and 4.5 mm ventral from bregma. The Isoflurane dosage was then adjusted to 2.5 lpm. The needle was inserted into the right mPFC, and the injection was started. The injection was performed at a rate of 0.2 μL/min for 15 min. In total, 3 μL of a compound was injected. After the infusion was completed, the needle was left in place for 10 min and gradually ejected. Three mL of 0.9% NaCl were injected subcutaneously. The incision was closed with skin clips. Animals were placed into the heated recovery chamber and monitored for 2 h before returning to their home cages.

### Perfusion

Animals were anesthetized using Isoflurane and injected intraperitoneally with 0.9 ml of Euthasol (150 mg/kg; Virbac, Fort Worth, TX). After terminal anesthesia, the rats were perfused transcardially using the Perfusion Two™ system (Leica Biosystems, Chicago, IL). As recommended by the manufacturer, ice-cold 9.25% sucrose solution in distilled deionized water was used as the prewash solution, followed by 4% PFA. The brains were harvested and post-fixed overnight in 4% PFA. Subsequently, the brains were dehydrated with 30% sucrose solution in PBS at 4°C and allowed to sink entirely to the bottom of the container. After the dehydration, brains were embedded into Tissue-Tek optimal cutting temperature compound (OCT) on dry ice and stored at -80°C for cryosectioning.

### Cryosectioning

Before cryosectioning, the brains were equilibrated at -20°C in a cryostat (Leica CM3050 S, Leica Biosystems, Wetzlar, Germany). Subsequently, the brains were coronally sectioned at 10 μm thickness. The sections were mounted on slides and air dried for 60 min at -20°C for RNAscope.

### RNAscope

The RNAscope Multiplex Fluorescent v2 Assay was used to visualize mRNA expression (Advanced Cell Diagnostics, ACD; Newark, CA). This assay allows the simultaneous detection of up to four mRNA targets. RNAscope target probes TACE/ADAM17 (Cat# 1052461-C1, Advanced Cell Diagnostics, Inc.), DRD1 (Cat# 317031-C2, Advanced Cell Diagnostics, Inc.), and AIF-1 (Cat# 457731-C3, Advanced Cell Diagnostics, Inc.) were assigned probe channels: C1, C2, and C3, respectively. First, the slides with brain sections were washed in Phosphate-buffered saline (PBS) for 5 min at room temperature to remove the OCT and subsequently baked for 30 min at 60°C. Afterward, the slides were post-fixed by immersing them in prechilled 4% paraformaldehyde (PFA) in PBS for 15 min at 4°C.

Subsequently, the brain sections were dehydrated by immersing them in a series of ethanol solutions, with 50% for the first immersion, 70% for the second immersion, and 100% for the third and fourth immersions. Each immersion lasted for 5 min at room temperature. After the dehydration stage, the slides were air-dried for 5 min at room temperature. Hydrogen peroxide was added and incubated for 10 min at room temperature. After incubation, the slides were washed with distilled deionized water (DDW) for 30 s at room temperature. The last step was repeated with fresh DDW. Subsequently, a target retrieval was performed by immersing the slides first in a boiling DDW for 10 seconds for acclimation and afterward in a boiling 1x Target Retrieval Reagent for 5 min. Next, the slides were washed in DDW for 15 s at room temperature with subsequent immersion in a 100% concentration ethanol for 3 min. Then, the slides were air-dried for 5 min at room temperature.

The hydrophobic barrier pen created a barrier around each brain section on a slide. Next, Protease III was added to each section, and the slides were incubated for 30 min at 40°C. After incubation, the slides were washed with DDW for 2 min at room temperature. The last step was repeated with fresh DDW. For hybridization, the TACE/ADAM17, DRD1, and AIF-1 probes were added to each slide and incubated for 2 hours at 40°C. After hybridization, the slides were washed in a washing buffer for 2 min at room temperature. The last step was repeated with a fresh washing buffer. The slides were immersed in 5x Saline Sodium Citrate and stored overnight at 4°C. Next, the slides were washed in a washing buffer for 2 min at room temperature. The last step was repeated with a fresh washing buffer, after which the amplification stage was launched. Amp 1 solution was added to each slide and hybridized for 30 min at 40°C. Afterward, the slides were washed in a washing buffer for 2 min at room temperature. The last step was repeated with a fresh washing buffer. The amplification step was repeated with Amp 2 and then with Amp 3 solutions. After the amplification stage had been completed, the channel development was started. The signal from channel 1 was developed by adding HRP-C1 solution to each slide and incubated for 15 min at 40°C. The slides were then washed in a washing buffer for 2 min at room temperature. The last step was repeated with a fresh washing buffer. Opal™ 520 dye (FP1487001KT, Akoya Biosciences) was assigned to channel 1 and added to each slide with subsequent incubation for 30 min at 40°C. Afterward, the slides were washed in a washing buffer for 2 min at room temperature. The last step was repeated with a fresh washing buffer. Next, an HRP blocker was added to each slide and incubated for 15 min at 40°C. All 40°C incubations used a humidity control chamber (HybEZ oven, ACDbio). The slides were washed in a washing buffer for 2 min at room temperature. The last step was repeated with a fresh washing buffer. The signals from channels 2 and 3 were developed similarly, and Opal™ 570 dye (FP1488001KT, Akoya Biosciences) and Opal™ 620 dye (FP1495001KT, Akoya Biosciences) were assigned to channels 2 and 3, respectively. Finally, the slides were counterstained with DAPI and stored at 4°C for microscopy.

### Confocal Microscopy

All the slides were scanned with the Zeiss LSM 710 NLO confocal microscope (Zeiss, White Plains, NY). Wavelength absorbance-emission values were as follows: DAPI (410-449); ADAM17 (484-552); Drd1 (552-601); Aif (599-670). Using an oil immersion objective, z-stacks of the mPFC were obtained at 63x magnification. Additional images were captured using an Andor BC43 Spinning Disk Confocal system (Andor Technology, Belfast) using an oil immersion objective (Plan Apo 60x, NA 1.4; Nikon). For excitation, 405 nm, 488 nm, 561 nm, and 640nm lasers were used in sequence. Emission light was detected by an Andor sCMOS camera (4.2MP; 6.5 um pixel size).

### RNAScope Image Analyses

For image analysis, we used the HALO platform (Indica Labs, Albuquerque, NM) with multiplex fluorescence in-situ hybridization (FISH) module. Quantitative gene expression evaluation was performed at single-cell resolution. The multiplex FISH module allows quantifying RNA FISH probes on a cell-by-cell basis. Single cells were identified using nuclear dye DAPI, and the TACE/ADAM17, DRD1, and AIF-1 probes were measured within the cell membrane and presented as spots per cell.

### Automated Observation Cage Behavioral Measures

We used the PhenoTyper cages for behavioral assessment instrumented observation (Noldus Information Technology BV, Wageningen, the Netherlands). Instrumented observation cages consist of a bottom plate that represents a black square arena; four replaceable transparent walls with ventilation holes at the top; an illuminated shelter that can be controlled with a hardware control module to switch the light automatically on when the animal enters the shelter, or choose a specific shelter entrance; a top unit that contains an infrared sensitive camera with three arrays of infrared light-emitting diode (LED) lights, and a range of sensors and stimuli, including adjustable light conditions to create a day/night cycle, the single tone for operant conditioning test, or the white spotlight for approach-avoidance behavior testing. Instrumented observation cages allow testing of different behavioral characteristics of laboratory rodents in an environment like their home cage. Dark bedding was used to facilitate detection. The rats were observed in the instrumented observation cages for 48 h, including 12 h of acclimation. The entire observation period was recorded and analyzed using EthoVision XT video tracking software, including the Rat Behavior Recognition module (Noldus Information Technology). We measured grooming, jumping, supported rearing, unsupported rearing, twitching, sniffing, walking, resting, eating, and drinking. In addition, we measured eating frequency and duration using feeding monitors incorporating a beam break device (Noldus).

### Acoustic startle reflex (ASR)

We performed the ASR experiments during the light phase using the SR-LAB acoustic chambers (San Diego Instruments, San Diego, CA, USA). The rats were placed inside the Plexiglas startle enclosures that contained piezoelectric transducers and motion sensors to measure the startle magnitudes. We performed calibration of the acoustic stimuli intensities and response sensitivities before initiating the experiments. The experimental sessions started with a 5-min acclimation period with the following background noise and light conditions: background noise = 55 decibels (dB); light conditions = 400 lx, which were maintained during the whole session. After the acclimation period, the rats were exposed to a series of 30 tones, maintaining a 30-s interval between trials, with 10 tones at each intensity: 90 dB, 95 dB, and 105 dB. The duration of each acoustic stimulus was 20 milliseconds (ms) with a quasi-random order of trial exposures. Accelerometer readings were obtained at 1 ms intervals for 200 ms after the startle-inducing acoustic stimulus. The overall duration of the experiment was 22 min. All the measurements were recorded using SR-LAB startle software. The ASR results were normalized to body weight and averaged.

### Social Y-Maze (SYM)

Sociability was assessed during the light phase using a modified Y maze and implementing a protocol adapted from Vuillermot et al., 2017.^37^ The test was previously used to measure rodent social interactions.^38^ The Social Y-Maze (Conduct Science, Maze Engineers, Skokie, IL) is a plexiglass Y-maze with a triangular center (8 cm sides) and three identical arms (50 cm x 9 cm x 10 cm, length x width x height). One arm functions as the start arm, while the other two are equipped with rectangular wire mesh cages to hold the live conspecific or a ‘dummy object.’ The protocol was conducted using one social interaction test trial without habituation training. The animal was placed into the start arm and allowed to explore the maze for 9-min freely. An unfamiliar conspecific (same sex as the test animal) was put into one of the rectangular wire mesh cages. At the same time, a ‘dummy object’ made of multicolored LEGO® pieces (Billund, Denmark) was placed into the other wire mesh cage. The conspecifics and ‘dummy objects’ were counterbalanced across arms and treatment groups. The maze was cleaned with 70% ethanol and allowed to dry between trials. A camera was mounted directly above the maze to record each test session. Videos recordings were analyzed using Ethovision XT tracking software (Noldus Information Technology).

### Statistical Methods

We analyzed the data using GraphPad’s Prism version 9.0. Shapiro-Wilk statistical analyses were used to determine sample distribution. The Brown-Forsyth test was used to test for the equality of group variances. Two-way analysis of variance (ANOVA) was used when appropriate to examine the effect of the intervention, stress, and interaction between factors on outcome measures. Multiple comparisons were made using Tukey’s test. The ROUT method was used to investigate outliers. Differences were considered significant when *p*<0.05. The data is shown as the mean ± standard error of the mean.

## RESULTS

The tumor necrosis factor alpha-converting enzyme / a disintegrin and metalloproteinase 17 (TACE/ADAM17) is critical for the cleavage of different growth factors and inflammatory mediators.^22^ We reported a model in which supraoptimal TACE/ADAM17 activities may contribute to neuroinflammation and altered brain maturation.^26^ This follow-up study tests the hypothesis that TACE/ADAM17 contributes to behavioral alterations associated with stress-induced obesity. Male and female Lewis rats underwent experimental manipulations and behavioral readouts during critical brain maturational periods (**Figure 1**).

To examine the functional significance of TACE/ADAM17 in the stressed rat mPFC (**Figure 2A**), we administered a rat sequence-specific siRNA to attenuate TACE/ADAM17 mRNA levels. First, we performed a non-targeting control siRNA injection to validate siRNA diffusion into mPFC cells. Intracerebral injection of non-targeting control siRNA allows for assessing the siRNA delivery and uptake by the brain cells. The non-targeting siRNA is labeled with 6-FAM and can be visualized with fluorescence microscopy. Our results indicate efficient delivery and uptake of siRNA by the PFC brain cells (**Figure 2B1**).

**Figure 2.**
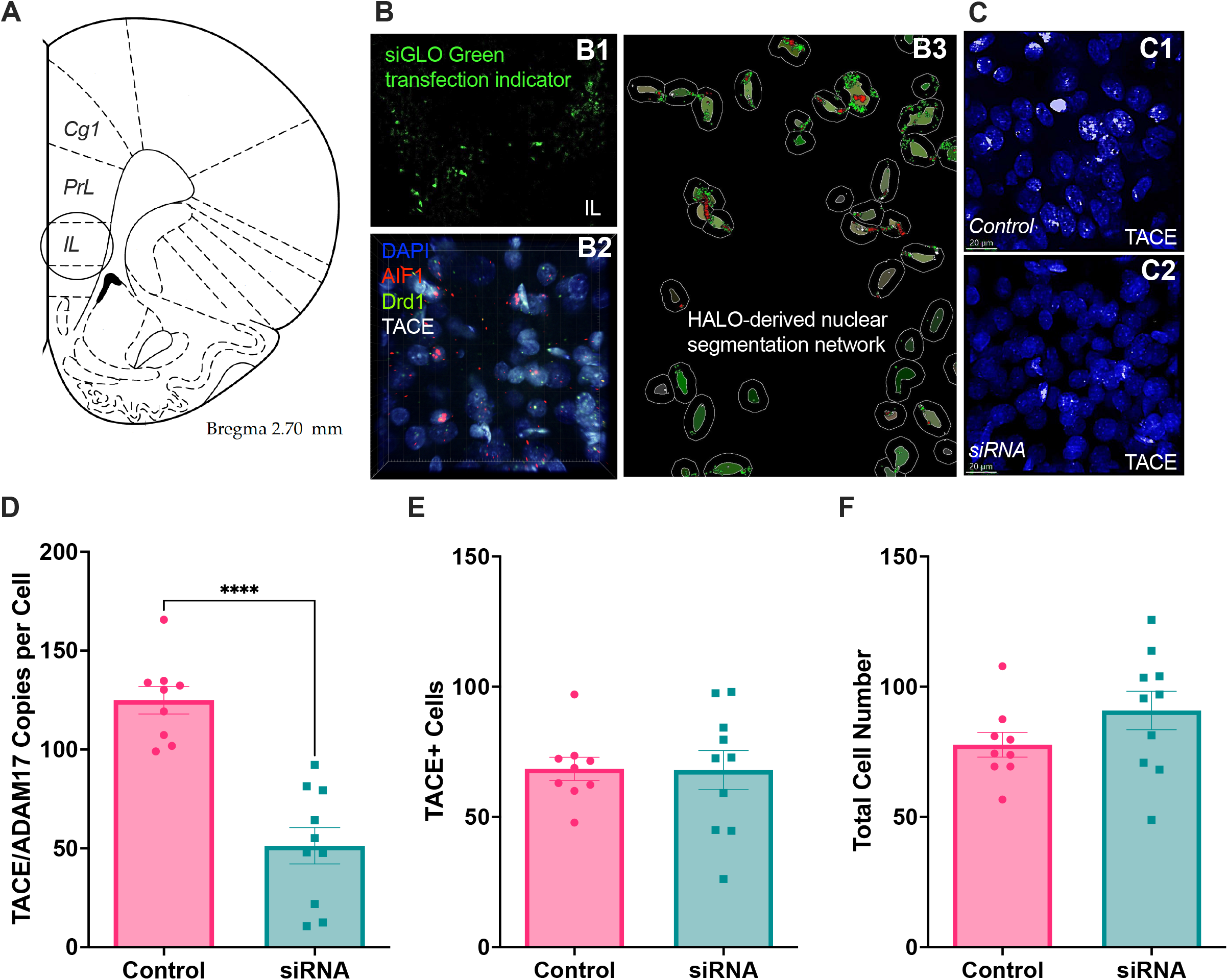
Intracerebral injection of TACE/ADAM17 siRNA significantly decreased TACE/ADAM17 mRNA levels in the mPFC. **(A)** Illustration from Paxinos and Watson rat brain atlas depicting mPFC injection site (cg1, cingulate cortex area; PrL, prelimbic cortex; infralimbic cortex; IL). **(B1)** Photomicrograph of rat IL injected with siGlo oligos, showing siRNA diffusion. **(B2)** Representative photomicrograph of merged RNAScope z-stacks performed to determine TACE/ADAM17, AIF1, and DRD1 mRNA levels in the mPFC. **(B3)** The HALO platform with multiplex fluorescence in-situ hybridization (FISH) module was used for nuclear segmentation, quantification, and analyses. Representative photomicrographs of vehicle control **(C1)** and siRNA **(C2)** injected brains showing reduced TACE/ADAM17 mRNA levels in the mPFC of siRNA-treated rats. **(D)** Analyses confirmed that the TACE/ADAM17 siRNA administration significantly reduced TACE/ADAM17 mRNA. **(E)** TACE/ADAM17 siRNA administration did not alter the total number of TACE/ADAM17 positive cells in the mPFC. **(F)** TACE/ADAM17 siRNA injections did not alter the total cell number in the mPFC. Scale bars = 20 micrometers. Controls, n = 9 rat brains; siRNA = 10 rat brains. ****, *p*<0.0001.

Next, we performed intracerebral injection of siRNA and measured TACE/ADAM17 mRNA levels with a fluorescent RNA in situ hybridization method (**Figure 2 B2-3**). As expected, TACE/ADAM17 siRNA significantly decreased TACE/ADAM17 mRNA levels in the brain compared to the vehicle-treated group at 72 h post-injection (*t*_17_=6.28, *p*<0.0001) (**Figure 2C-D**). This finding confirms the efficiency of the siRNA to silence TACE/ADAM17 mRNA levels. The number of TACE/ADAM17-positive cells did not change in the TACE/ADAM17 siRNA-treated group compared to the vehicle-treated group (*t*_17_=0.055, *p*=0.96) (**Figure 2E**). Similarly, the total cell number also was not affected by the TACE/ADAM17 siRNA administration (*t*_17_=1.46, *p*=0.16) (**Figure 2F**). These data demonstrate that the TACE/ADAM17 siRNA was not toxic to cells.

### mPFC TACE/ADAM17 siRNA influences behavioral profiles

Having established the efficacy of the intracerebral siRNA injection to attenuate TACE/ADAM17, we decided to examine whether the intervention ameliorated behavioral proxies related to prefrontal network integrity (**Supplemental Figure 1**). Considering the fundamental roles of prefrontal networks on top-down behavior control, we next monitored the TACE/ADAM17 siRNA injection effects on ethologically relevant behaviors (**Table 1**). Optimal gene knockdown at the mRNA level is usually reached at 48-72 h after Accell™ siRNA delivery; thus, we commenced to measure behavioral outcomes after this period. We used Noldus’ PhenoTyper home cages and Rat Automated Behavior Recognition (ABR) module to measure ten relevant behaviors for 48 h (**Figure 3A**). Monitoring behaviors in this naturalistic environment over long periods provides complex and ethologically relevant behaviors that reflect how interventions impact conserved endophenotypes. We identified distinct behavioral profiles (**Figure 3B-C**) (behavior: F_9, 530_=986.20, *p*<0.0001; group: F_5, 530_=0.033, *p*=1.00; interaction: F_45, 530_=6.37, *p*<0.0001). TACE/ADAM17 siRNA delivery to the mPFC reduced eating behavior (post hoc *p*=0.013) and increased grooming (post hoc *p*=0.007) in stressed rats relative to stressed rats receiving control injections (**Figure 3C**). Analyses of total ambulatory behavior in the PhenoTyper cages revealed a significant time and interaction effect (time: F_4.01, 189.1_=26.38, *p*<0.0001; group: F_5, 53_=1.52, *p*=1.52; interaction: F_110, 1038_=1.28, *p*=0.031). TACE/ADAM17 siRNA administration reduced ambulation relative to control stress-exposed rats at 04:00 h (*p*=0.011) (**Figure 3D**).

**Figure 3.**
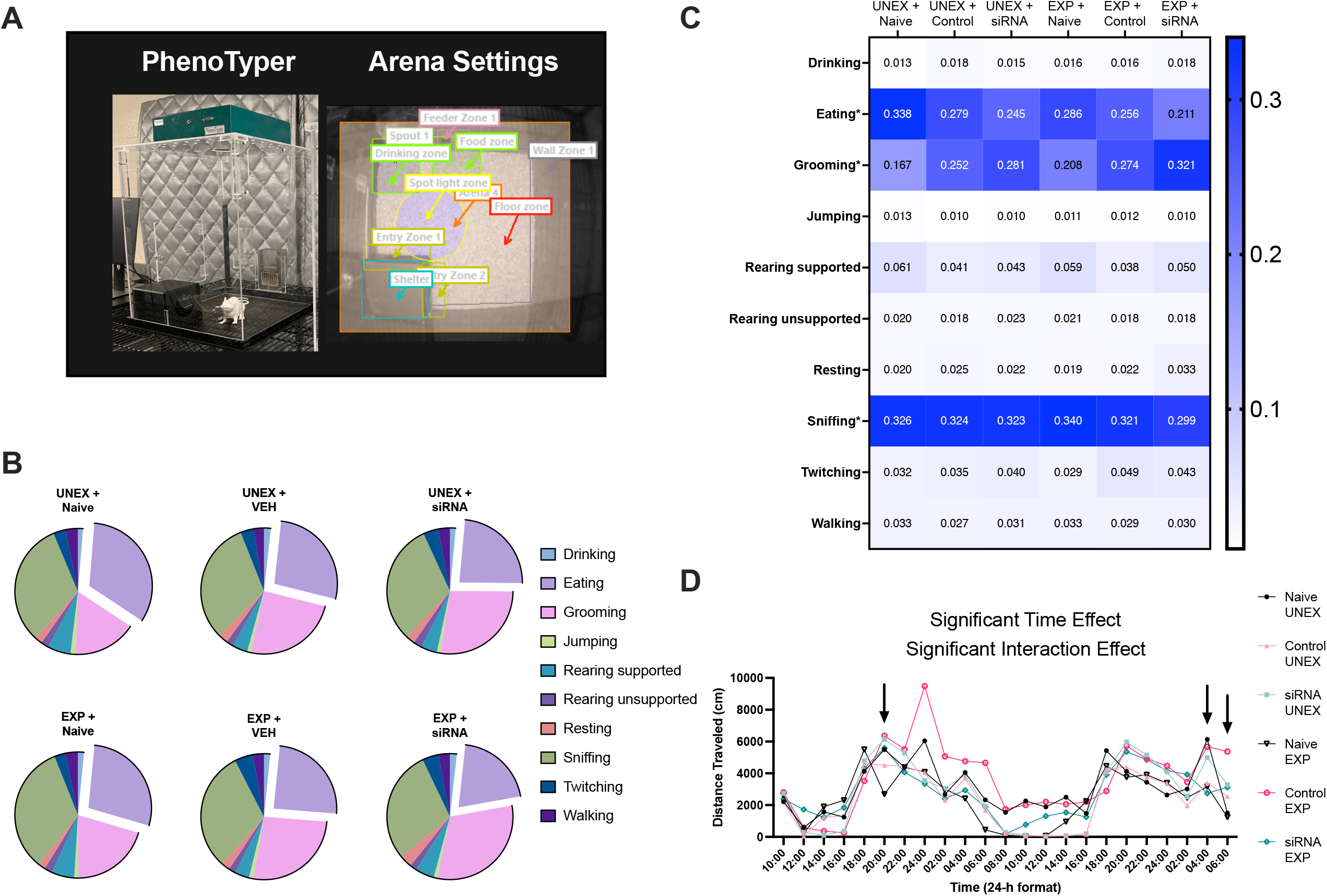
mPFC TACE/ADAM17 siRNA administration influences grooming, eating, and ambulation behaviors in stress-exposed rats. **(A)** Home cage monitoring apparatus and arena settings illustrating relevant zones. **(B-C)** Exploding pie charts and heatmap illustrating behavioral profiles from the rats that underwent home cage behavioral monitoring. Automated quantification of 10 behavioral parameters for two days demonstrates significant differences between groups’ behavioral profiles. In stressed rats, TACE/ADAM17 siRNA administration reduced the behavioral probability of eating (difference: 0.044; 95% CI of difference: 0.0060 to 0.083) and increased the behavioral probability of grooming (difference: -0.047; 95% CI of difference: -0.085 to - 0.0082) relative to controls. **(D)** Total distance traveled analyses revealed a significant time and interaction effect. Rats exposed to stress and receiving the TACE/ADAM17 siRNA injections exhibited reduced ambulation relative to control stress-exposed animals (at 04:00 h: difference: -2922; 95% CI of difference: -5198 to -645.80). Naïve Unexposed, n = 8; Naïve Exposed, n = 8; Control Unexposed, n = 12; Control Exposed, n = 8; siRNA Unexposed, n = 10; siRNA Exposed, n = 13. Asterisks (*) denote behaviors significantly affected by the experimental conditions based on two-way ANOVA and post hoc analyses (grooming, eating, and sniffing).

Food intake is a fundamental behavior that is often altered following stress. We tested this notion in our model and found that exposure to adolescent traumatic stress and social isolation increased eating behaviors (**Supplemental Figure 1B-C**). Eating frequency analyses showed significant time and interaction effects (time: F_8.79, 324.7_=19.18, *p*<0.0001, group: F_5, 53_=0.46, *p*=0.81; interaction: F_110, 813_=1.65, *p*<0.0001) (**Figure 4A**). Stressed rats receiving the control vehicle injection exhibited a shift in their eating pattern, with a reduced eating frequency during the light cycle. We found that the stressed rats that received the TACE/ADAM17 siRNA injection exhibited increased eating frequency relative to control stress-exposed animals at 18:00 h on both days tested (*for day 1*: *p*=0.022; *for day 2*: *p*=0.0091) (**Figure 4A**), approximating the eating frequency of unexposed rats. Eating duration analyses revealed significant time and group effects (time: F_9.92, 367.3_=21.32, *p*<0.0001; group: F_5, 53_=3.75, *p*=0.0056; interaction: F_110, 815_=1.11, *p*=0.22), with surgically naïve rats spending more time eating relative to rats receiving the intracerebral injections (**Figure 4B**). Cumulative eating duration measures confirmed this finding, demonstrating that the rats that underwent surgeries spent less time eating (treatment: F_2, 49_=11.29, *p*<0.0001; stress: F_1, 49_=0.53, *p*=0.47; interaction: F_2, 49_=2.91, *p*<0.0.063) (**Figure 4C**). Interestingly, TACE/ADAM17 siRNA increased the number of feeding bouts relative to vehicle controls. The treatment showed opposite effects based on stress exposure (treatment effect: F_2, 49_=3.30, *p*=0.045; stress: F_1, 49_=0.17, *p*=0.68; interaction effect: F_2, 49_=3.47, *p*=0.039; Tukey’s post hoc for unexposed groups: *p*=0.017) (**Figure 4D**). Heatmaps demonstrated increased duration in the feeding zone in stress-exposed rats (14.4 and 12.5%) relative to vehicle controls (10.3 and 10.8%) (**Figure 4E**). Notably, TACE/ADAM17 siRNA-treated animals spent less time in the feeding zone than stress-exposed vehicle controls (12.5 vs. 14.4% for stressed rats). While the surgical intervention altered some behavioral outcomes (relative to surgically naïve rats), the siRNA and vehicle-treated rats displayed similar acoustic startle reflexes (ASR) at one-week post-surgery (**Supplemental Figure 2**).

**Figure 4.**
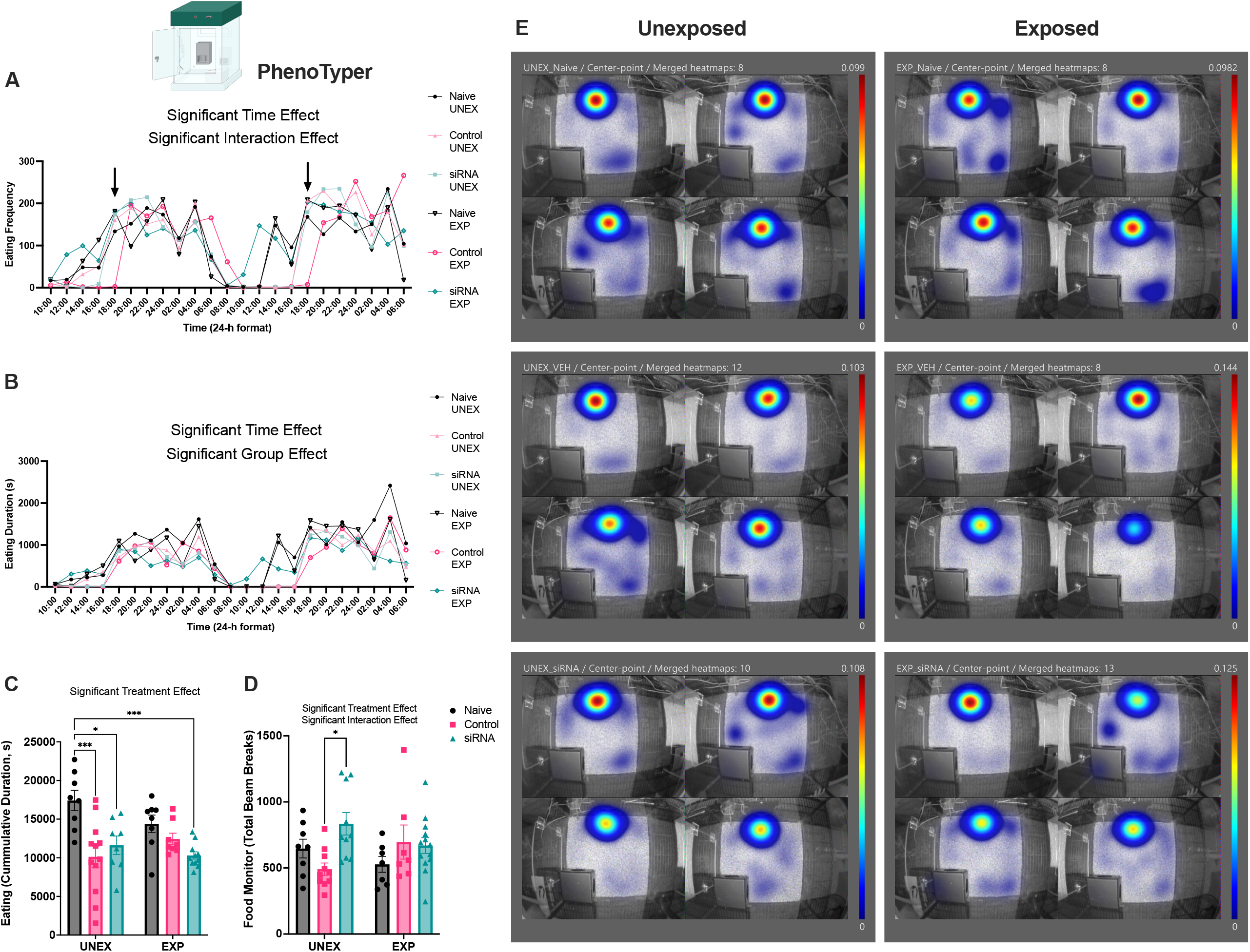
TACE/ADAM17 siRNA injection to the mPFC influences feeding frequency and duration. **(A)** Eating frequency data based on the rat automated behavior recognition module. Stressed rats receiving the control injection exhibited reduced eating frequency during the day. Rats exposed to stress and receiving the TACE/ADAM17 siRNA injections exhibited increased eating frequency relative to control stress-exposed animals at 18:00 h on both days tested (arrows: *for day 1*: *p*=0.022, difference: 175.9, 95% CI of difference: 25.13 to 326.70; *for day 2*: *p*=0.0091, difference: 194.70, 95% CI of difference: 47.72 to 341.70). **(B)** Eating duration data based on the rat automated behavior recognition module. In general, surgically naïve rats exhibited longer eating durations than those that underwent surgery. **(C)** The cumulative eating duration was reduced in the rats that underwent surgery relative to naïve controls. **(D)** Food monitor data showed that unexposed rats receiving the TACE/ADAM17 siRNA infusion exhibited more feeding bouts than vehicle controls over the 48-h testing period. **(E)** Representative *mean* merged heatmaps illustrating average distribution in the feeding zone for each group. The maximum is expressed as a fraction of the time in the feeding zone (pixel color denotes the average proportion of a track found at the feeding zone). Please note that when rats keep moving across the arena, this value can be low, for example, 0.01. This means that the region of peak occurrence contained 1% of the positions. Heatmaps show increased duration in the feeding zone in surgically manipulated and stress-exposed rats (14.4 and 12.5%) relative to vehicle controls (10.3 and 10.8%). Stress-exposed and TACE/ADAM17 siRNA-treated animals spent less time in the feeding zone than exposed vehicle controls (12.5 vs. 14.4%). Naïve Unexposed, n = 8; Naïve Exposed, n = 8; Control Vehicle (VEH) Unexposed, n = 12; Control Vehicle (VEH) Exposed, n = 8; siRNA Unexposed, n = 10; siRNA Exposed, n = 13. *, *p*<0.05; ***, *p*<0.001.

### Long-term effects of TACE/ADAM17 siRNA administration on the acoustic startle and eating and social behaviors

We planned a secondary set of experiments to evaluate the long-term effects of prefrontal TACE/ADAM17 siRNA administration on stress-relevant behaviors (**Figure 5A**). The cumulative eating duration was similar between groups (stress: F_1, 71_=0.035, *p*=0.85; treatment: F_2, 71_=1.71, *p*=0.19; interaction: F_2, 71_=0.60, *p*=0.55) (**Figure 5B**). We examined binge eating-like behaviors in a subcohort of animals. We found that TACE/ADAM17 siRNA rats that were given intermittent access to an obesogenic diet consumed more food than vehicle controls at 2.5 h after reintroducing the obesogenic diet (**Supplemental Figure 3**). The rats exposed to traumatic stress and surgical manipulations exhibited blunted ASR responses (stress: F_1, 64_=8.70, *p*=0.0045; treatment: F_2, 64_=3.56, *p*=0.034; interaction: F_2, 64_=2.58, *p*=0.084) (**Figure 5C**). This effect was not associated with changes in the latency to respond to the acoustic stimuli (stress: F_1, 71_=1.09, *p*=0.30; treatment: F_2, 71_=0.65, *p*=0.52; interaction: F_2, 71_=0.39, *p*=0.68) (**Figure 5D**). We found that stress and TACE/ADAM17 siRNA administration did not significantly impact either duration with conspecific (interaction: F_2, 70_=0.84, *p*=0.43; stress: F_1, 70_=0.28, *p*=0.60; treatment: F_2, 70_=0.44, *p*=0.65) (**Figure 5E**) or distance traveled (interaction: F_2, 72_=1.31, *p*=0.28; stress: F_1, 72_=0.00054, *p*=0.98; treatment: F_2, 72_=1.16, *p*=0.32) (**Figure 5F**).

**Figure 5.**
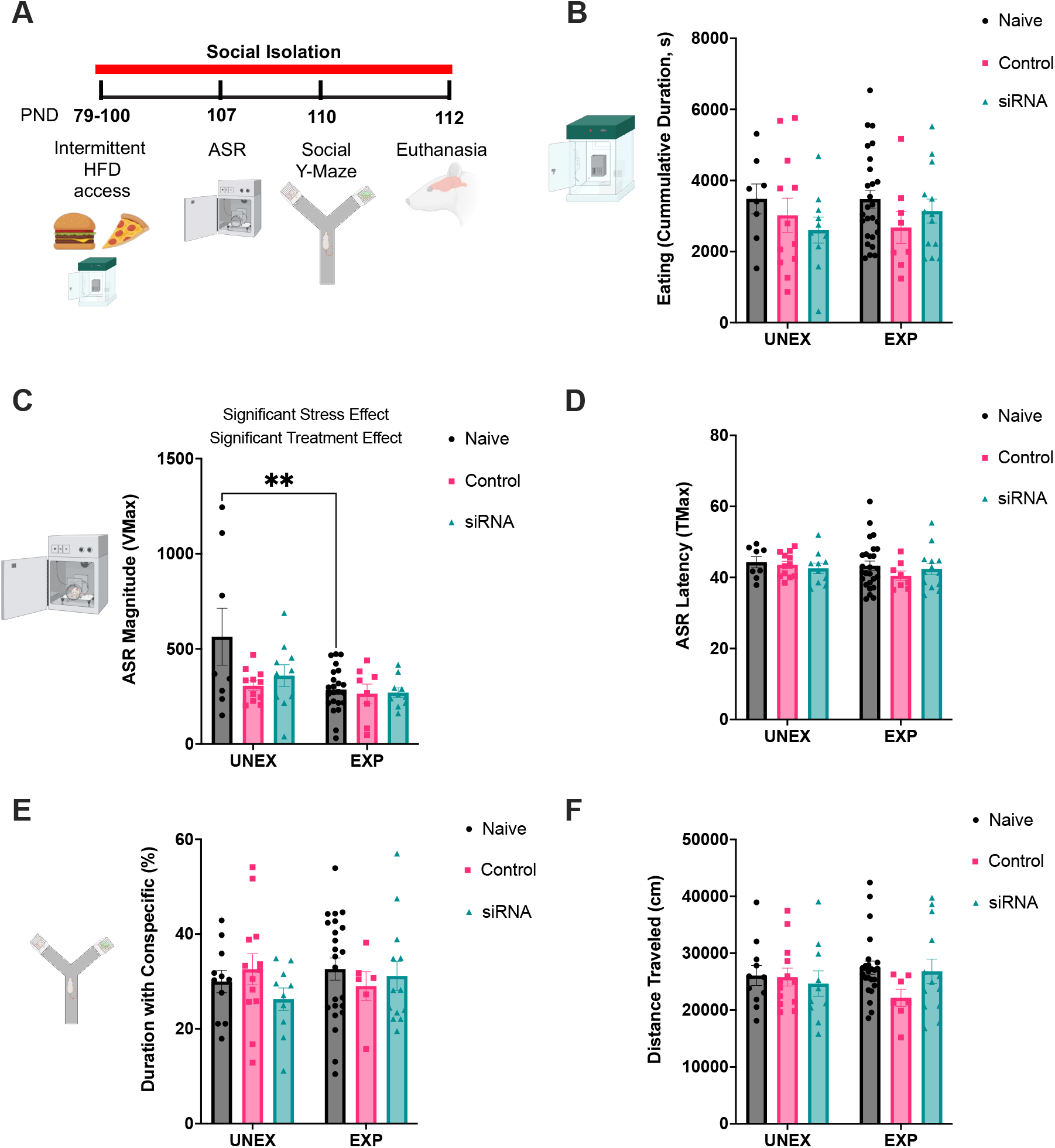
Effects of TACE/ADAM17 siRNA administration to the mPFC on social behavior and acoustic startle. **(A)** Study design and timeline of experimental procedures, behavioral tests, and outcome measures to examine the long-term effects of TACE/ADAM17 siRNA mPFC administration in a rat model of PTSD. **(B)** The experimental conditions did not significantly affect the cumulative eating duration in the PhenoTyper cages. **(C)** Adolescent trauma attenuated the magnitude of the ASR. Further, the rats undergoing surgical manipulations exhibited blunted startle reactivity relative to surgically naïve rats. **(D)** ASR attenuation was not associated with changes in the latency to respond to the acoustic stimuli. **(E-F)** Experimental conditions did not alter social behaviors in a social Y maze. **(E)** The time spent interacting with same-sex conspecific was similar between groups (expressed as a percentage of the total time; 9 min test). **(F)** The experimental conditions did not affect the total distance traveled in the social Y maze. Naïve Unexposed, n = 8; Naïve Exposed, n = 8; Control Vehicle Unexposed, n = 12; Control Vehicle Exposed, n = 8; siRNA Unexposed, n = 10; siRNA Exposed, n = 13. **, *p*<0.01.

### TACE/ADAM17 siRNA targets prefrontal cortical microglia

TACE/ADAM17 plays a critical role in neuroinflammatory responses by cleavage and release of critical proinflammatory mediators.^39^ We examined the TACE/ADAM17 siRNA effects on a critical marker of neuroinflammation. The allograft inflammatory factor 1 (AIF1) is a typical marker for microglia and colocalizes with TACE/ADAM17 (**Figure 6A**). We found that knocking down TACE/ADAM17 mRNA significantly decreased the AIF1 expression level in the mPFC at 72 h post-injection (*t*_17_=8.78, *p*<0.0001) (**Figure 6B**). The percentage of AIF+ cells that expressed TACE/ADAM17 was also reduced in the mPFC of siRNA-treated rats relative to controls (*t*_17_=3.50, *p*=0.0027) (**Figure 6C**).

**Figure 6.**
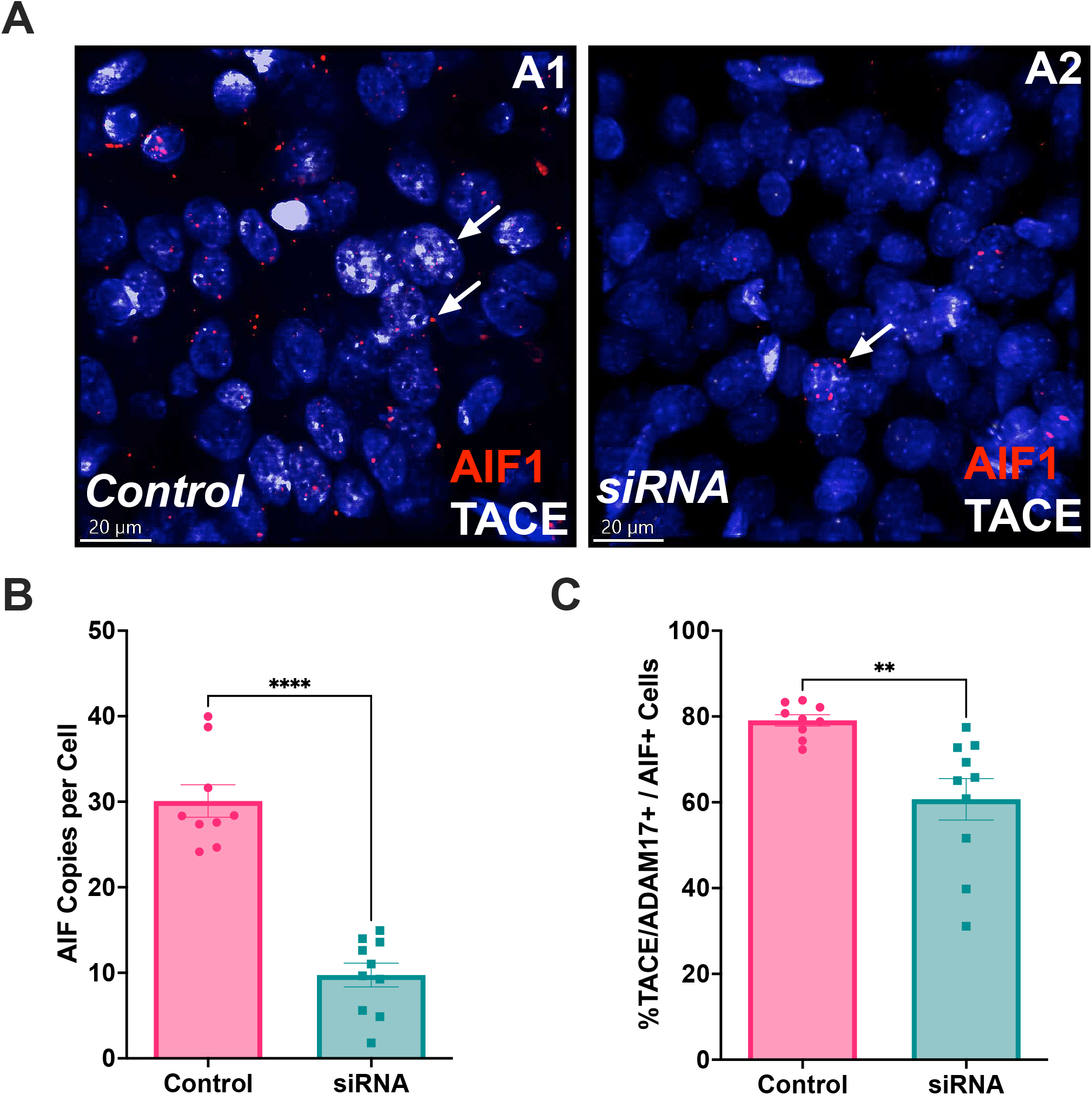
TACE/ADAM17 siRNA intracerebral injection attenuates the expression of a crucial neuroinflammation biomarker in the mPFC. **(A)** Representative RNAScope photomicrographs demonstrating allograft inflammatory factor 1 (AIF1; also known as ionized calcium-binding adapter molecule 1 or Iba-1) expressing cells that co-express TACE/ADAM17 mRNA. Representative sections from control-treated rat brain **(A1)** and TACE/ADAM17 siRNA-treated rat brain **(A2)** show decreased TACE/ADAM17 and AIF mRNA levels in siRNA-treated animals. **(B)** TACE/ADAM17 siRNA significantly decreased the AIF1 mRNA levels in the mPFC. **(C)** The percentage of AIF+ cells that expressed TACE/ADAM17 mRNA was also reduced in the mPFC of siRNA-treated rats. Scale bars = 20 micrometers. Controls, n = 9 rat brains; siRNA = 10 rat brains. **, *p*<0.01; ****, *p*<0.0001.

## DISCUSSION

This study investigated the effects of adolescent traumatic stress on multiple behavioral domains. We examined the involvement of mPFC TACE/ADAM17 on the startle reflex, social behavior, home cage naturalistic behaviors, and neuroinflammation. We found that adolescent traumatic stress blunts startle reactivity and alter eating behaviors (increased intake and disrupted eating patterns) in a rat model of posttraumatic stress disorder (PTSD). We also found that the rats that received TACE/ADAM17 siRNA administration in the medial prefrontal cortex (mPFC) exhibited decreased eating behaviors compared to controls. These changes were associated with decreased AIF-1 expression in the mPFC (a typical marker of microglia and neuroinflammation).

The formation of connections in mPFC takes place during prenatal development and continues postnatally. This process is significantly shaped by synaptogenesis and synaptic pruning, which occur concurrently in a balanced manner. Adolescence is considered a very important development period because of the active organization of neuronal formation, particularly in the prefrontal cortex. Formational processes in the prefrontal cortex are disturbed by environmental factors, including stress, which leads to aberrant development of neuronal circuitry in the PFC, resulting in specific phenotypes that may continue to adulthood.^40^

Stress and PTSD are considered risk factors for obesity. Stress is also recognized as one of the primary triggers for binge eating. Furthermore, these factors present an increased risk for metabolic syndrome development.^41,42^ We reported that exposure to an obesogenic environment during adolescence alters the structural integrity and maturation of the mPFC.^32^ Feeding patterns are complex behavioral expressions constructed with different areas of the brain and circuits connecting important centers involved in the formation of feeding behavior, including top-down control mechanisms.^43^

The mPFC plays a central role in the feeding pattern regulatory system, providing top-down control of crosstalk between circuits involved in different aspects of feeding behavior. Impulses from the mPFC are transmitted to the basolateral amygdala and central nucleus of the amygdala, the bed nucleus of stria terminalis, and the nucleus accumbens. Subsequently, these impulses are transmitted to the lateral hypothalamic area, ventral trigeminal area, and other regions. These projections that integrate the crosstalk between the mPFC and diencephalic centers of satiety and hunger are essential in controlling food intake and characterizing feeding patterns.^44-47^ Satiety and hunger centers are in the hypothalamus and are integral to the feeding pattern construct. These centers connect with different brain areas, including the frontal lobe, which processes nutritional requirements and metabolic homeostasis information. Hypothalamic regions implicated in satiety and hunger are also heavily connected to the immune and endocrine systems, which control different aspects of feeding behavior.^48-51^ Metalloproteases play a critical role in CNS maturation and function, providing crucial proteolytic shedding activities.^52-54^ During CNS development, a disintegrin and metalloproteinases (ADAMs) significantly influence neuronal differentiation, proliferation, migration, and axonal myelination.^52^ TACE/ADAM17 has a critical functional role in neuronal development, including activation of neural cell adhesion and neurite outgrowth.^55-57^ It has been shown that TACE/ADAM17 regulates amyloid precursor protein (APP), which is assumed to play a role in neuronal migration and synaptic connectivity during development.^58-60^ Some studies indicate that TACE/ADAM17 is involved in producing secreted amyloid precursor protein alpha (sAPPα), a non-amyloidogenic fragment and soluble.^58^ TACE/ADAM17-dependent APP processing through non-amyloidogenic pathways might play a role in Alzheimer’s disease.^61,62^ TACE/ADAM17 is also involved in neuronal development, mainly regulating the signaling pathway through the epidermal growth factor receptor (EGF-R).^53^ It has also been shown that EGF-R signaling plays a critical role in neuronal development and synaptic plasticity, and memory formation.^63,64^

Obesogenic environments create conditions for TACE/ADAM17 upregulation during adolescence. We demonstrated that consuming an obesogenic high-saturated fat diet during adolescence increased TACE/ADAM17 protein levels in the brain, specifically in the hippocampus.^26^ Similarly, consuming an obesogenic diet significantly increased pro-inflammatory mediators, particularly tumor necrosis factor-alpha (TNF-α), associated with robust neuroinflammatory states. These conditions were correlated with alterations in brain maturation and structure, including significant changes in volumetric parameters.^26^ TNF-α and its receptors are critical substrates for TACE/ADAM17, indicating the involvement of this protease in regulating inflammatory processes.^52,55,65^ This study demonstrated that TACE/ADAM17 silencing significantly reduced AIF-1 gene expression, a sensitive marker of microglia cell number and activities. Our results affirm that TACE/ADAM17 is crucial in neuroinflammatory signaling.^39^

Our study demonstrates a mechanistic connection between TACE/ADAM17 and microglia. Microglia are phagocytic scavenger cells and resident macrophages of the central nervous system (CNS) that, unlike other glial cells, do not originate from neuroectoderm but are derived from the mesoderm.^66,67^ These cells possess mobile processes and are distributed throughout the gray and white matter. Microglia can migrate within the central nervous system and scan the surrounding environment for potential harmful components, including microbes, serving as an immune defense mechanism in the CNS.^68-70^ One of the critical characteristics of microglia is that, when activated, these cells can secrete inflammatory mediators, like nitric oxide and glutamate, in response to tissue damage or microorganisms.^71-73^ Microglia play a critical role in neuronal expansion and differentiation, contributing to synaptic formation during development. Alterations in brain formation due to traumatic stress exposures during development may provoke microglia activation, which can significantly affect regulatory mechanisms involved in synaptogenesis and synaptic pruning processes and influence the course of maturation.^74,75^ Our results suggest that microglia activities in the PFC are regulated through pathways involving TACE/ADAM17.

Some of the aspects of this study represent limitations and require further assessment.

The spatiotemporal expression of mPFC TACE/ADAM17 and associated cytokine profiles must be determined to clarify a mechanism better. Our results do not clarify whether the TACE/ADAM17 silencing effects are mediated by microglia, a change in the extracellular milieu via intermediate factors, or in response to neuronal activities. Experimentation should be carried out to disentangle the relative contribution of neural phenotypes to the observed effects. TNF receptor inhibitors need to be considered to dissect the TACE/ADAM17-TNF signaling axis. The effect of a single siRNA intracerebral injection and multiple behavioral tests on the same animal may also confound the interpretation of the study. Future experiments with genetic or pharmacological tools that allow sustained blockade are required to dissociate the chronic vs. acute roles of TACE/ADAM17 in stress-induced neuroinflammation. While females were included in this study, critical experiments should be replicated, and statistical analyses performed to identify sex-specific differences. The results of this study require cautious interpretation, specifically when extrapolating to human conditions, since there are differences between rats and humans as it pertains to the PFC and pathophysiological processes related to stress-induced neuroinflammation and disordered eating.^76^ Studies accentuating other brain areas involved in trauma-induced neuroinflammation and obesity are required. Alternate silencing pharmacological approaches are warranted.

In summary, the current study validates the pathophysiological model we reported demonstrating that TACE/ADAM17 is involved in neuroinflammation and may play essential roles in regulating feeding patterns under obesogenic conditions. We report an interaction between TACE/ADAM17 and neuroinflammatory marker AIF-1 in the mPFC. We also demonstrated that mPFC TACE/ADAM17 influences feeding patterns in rats exposed to traumatic experiences during adolescence. Together, our study supports that TACE/ADAM17 represents a promising target to ameliorate the impact of adolescent traumatic stress on the brain and behavior.

## ACKNOWLEDGMENTS

This study was partly supported by the NIH (DK124727, GM060507, MD006988) and the Loma Linda University School of Medicine GRASP Seed Funds to JDF.

## DATA AVAILABILITY

In addition to the data presented in the supplementary materials, supportive datasets are available from the corresponding author upon reasonable request.

## REFERENCES

1. McLaughlin KA, Koenen KC, Hill ED, et al. Trauma exposure and posttraumatic stress disorder in a national sample of adolescents. J Am Acad Child Adolesc Psychiatry. 2013;52(8):815-830.e814.

2. Cuffe SP, Addy CL, Garrison CZ, et al. Prevalence of PTSD in a community sample of older adolescents. J Am Acad Child Adolesc Psychiatry. 1998;37(2):147–154.

3. Giaconia RM, Reinherz HZ, Silverman AB, Pakiz B, Frost AK, Cohen E. Traumas and posttraumatic stress disorder in a community population of older adolescents. J Am Acad Child Adolesc Psychiatry. 1995;34(10):1369–1380.

4. Jin H, Lanouette NM, Mudaliar S, et al. Association of posttraumatic stress disorder with increased prevalence of metabolic syndrome. J Clin Psychopharmacol. 2009;29(3):210–215.

5. Scott KM, Bruffaerts R, Simon GE, et al. Obesity and mental disorders in the general population: results from the world mental health surveys. Int J Obes (Lond). 2008;32(1):192–200.

6. Farr OM, Sloan DM, Keane TM, Mantzoros CS. Stress- and PTSD-associated obesity and metabolic dysfunction: a growing problem requiring further research and novel treatments. Metabolism. 2014;63(12):1463–1468.

7. Kanoski SE. Cognitive and neuronal systems underlying obesity. Physiol Behav. 2012;106(3):337–344.

8. Fuster JM. Frontal lobe and cognitive development. J Neurocytol. 2002;31(3-5):373–385.

9. Sowell ER, Peterson BS, Thompson PM, Welcome SE, Henkenius AL, Toga AW. Mapping cortical change across the human life span. Nat Neurosci. 2003;6(3):309–315.

10. Hodel AS. Rapid Infant Prefrontal Cortex Development and Sensitivity to Early Environmental Experience. Dev Rev. 2018;48:113–144.

11. Catani M. The anatomy of the human frontal lobe. Handb Clin Neurol. 2019;163:95–122.

12. Henri-Bhargava A, Stuss DT, Freedman M. Clinical Assessment of Prefrontal Lobe Functions. Continuum (Minneap Minn). 2018;24(3, behavioral neurology and psychiatry):704–726.

13. Xu P, Chen A, Li Y, Xing X, Lu H. Medial prefrontal cortex in neurological diseases. Physiol Genomics. 2019;51(9):432–442.

14. Rojkova K, Volle E, Urbanski M, Humbert F, Dell’Acqua F, Thiebaut de Schotten M. Atlasing the frontal lobe connections and their variability due to age and education: a spherical deconvolution tractography study. Brain Struct Funct. 2016;221(3):1751–1766.

15. Lowe CJ, Reichelt AC, Hall PA. The Prefrontal Cortex and Obesity: A Health Neuroscience Perspective. Trends Cogn Sci. 2019;23(4):349–361.

16. Caslini M, Bartoli F, Crocamo C, Dakanalis A, Clerici M, Carrà G. Disentangling the Association Between Child Abuse and Eating Disorders: A Systematic Review and Meta-Analysis. Psychosom Med. 2016;78(1):79–90.

17. Danese A, Tan M. Childhood maltreatment and obesity: systematic review and meta-analysis. Mol Psychiatry. 2014;19(5):544–554.

18. Imperatori C, Innamorati M, Lamis DA, et al. Childhood trauma in obese and overweight women with food addiction and clinical-level of binge eating. Child Abuse Negl. 2016;58:180–190.

19. Collin G, van den Heuvel MP. The ontogeny of the human connectome: development and dynamic changes of brain connectivity across the life span. Neuroscientist. 2013;19(6):616–628.

20. Mallya AP, Wang HD, Lee HNR, Deutch AY. Microglial Pruning of Synapses in the Prefrontal Cortex During Adolescence. Cereb Cortex. 2019;29(4):1634–1643.

21. Schalbetter SM, von Arx AS, Cruz-Ochoa N, et al. Adolescence is a sensitive period for prefrontal microglia to act on cognitive development. Sci Adv. 2022;8(9):eabi6672.

22. Sommer D, Corstjens I, Sanchez S, et al. ADAM17-deficiency on microglia but not on macrophages promotes phagocytosis and functional recovery after spinal cord injury. Brain Behav Immun. 2019;80:129–145.

23. Zhang S, Kojic L, Tsang M, et al. Distinct roles for metalloproteinases during traumatic brain injury. Neurochem Int. 2016;96:46–55.

24. Chemaly M, McGilligan V, Gibson M, et al. Role of tumour necrosis factor alpha converting enzyme (TACE/ADAM17) and associated proteins in coronary artery disease and cardiac events. Arch Cardiovasc Dis. 2017;110(12):700–711.

25. Vidal PM, Lemmens E, Avila A, et al. ADAM17 is a survival factor for microglial cells in vitro and in vivo after spinal cord injury in mice. Cell Death Dis. 2013;4(12):e954.

26. Vega-Torres JD, Ontiveros-Angel P, Terrones E, et al. Short-term exposure to an obesogenic diet during adolescence elicits anxiety-related behavior and neuroinflammation: modulatory effects of exogenous neuregulin-1. Transl Psychiatry. 2022;12(1):83.

27. Rose-John S. ADAM17, shedding, TACE as therapeutic targets. Pharmacol Res. 2013;71:19–22.

28. Yoon S, Baik JH. Dopamine D2 receptor-mediated epidermal growth factor receptor transactivation through a disintegrin and metalloprotease regulates dopaminergic neuron development via extracellular signal-related kinase activation. J Biol Chem. 2013;288(40):28435–28446.

29. Zunke F, Rose-John S. The shedding protease ADAM17: Physiology and pathophysiology. Biochim Biophys Acta Mol Cell Res. 2017;1864(11 Pt B):2059–2070.

30. Miller GE, Chen E, Zhou ES. If it goes up, must it come down? Chronic stress and the hypothalamic-pituitary-adrenocortical axis in humans. Psychol Bull. 2007;133(1):25–45.

31. Carpenter LL, Carvalho JP, Tyrka AR, et al. Decreased adrenocorticotropic hormone and cortisol responses to stress in healthy adults reporting significant childhood maltreatment. Biol Psychiatry. 2007;62(10):1080–1087.

32. Vega-Torres JD, Haddad E, Lee JB, et al. Exposure to an obesogenic diet during adolescence leads to abnormal maturation of neural and behavioral substrates underpinning fear and anxiety. Brain Behav Immun. 2018;70:96–117.

33. Cadoni C. Fischer 344 and Lewis Rat Strains as a Model of Genetic Vulnerability to Drug Addiction. Front Neurosci. 2016;10:13.

34. Kalyan-Masih P, Vega-Torres JD, Miles C, et al. Western High-Fat Diet Consumption during Adolescence Increases Susceptibility to Traumatic Stress while Selectively Disrupting Hippocampal and Ventricular Volumes. eNeuro. 2016;3(5).

35. Zoladz PR, Fleshner M, Diamond DM. Psychosocial animal model of PTSD produces a long-lasting traumatic memory, an increase in general anxiety and PTSD-like glucocorticoid abnormalities. Psychoneuroendocrinology. 2012;37(9):1531–1545.

36. Zoladz PR, Park CR, Fleshner M, Diamond DM. Psychosocial predator-based animal model of PTSD produces physiological and behavioral sequelae and a traumatic memory four months following stress onset. Physiol Behav. 2015;147:183–192.

37. Vuillermot S, Luan W, Meyer U, Eyles D. Vitamin D treatment during pregnancy prevents autism-related phenotypes in a mouse model of maternal immune activation. Mol Autism. 2017;8:9.

38. Weber-Stadlbauer U, Richetto J, Labouesse MA, Bohacek J, Mansuy IM, Meyer U. Transgenerational transmission and modification of pathological traits induced by prenatal immune activation. Mol Psychiatry. 2017;22(1):102–112.

39. Dominguez-Garcia S, Castro C, Geribaldi-Doldán N. ADAM17/TACE: a key molecule in brain injury regeneration. Neural Regen Res. 2019;14(8):1378–1379.

40. Shaw GA, Dupree JL, Neigh GN. Adolescent maturation of the prefrontal cortex: Role of stress and sex in shaping adult risk for compromise. Genes Brain Behav. 2020;19(3):e12626.

41. Razzoli M, Pearson C, Crow S, Bartolomucci A. Stress, overeating, and obesity: Insights from human studies and preclinical models. Neurosci Biobehav Rev. 2017;76(Pt A):154–162.

42. Gluck ME. Stress response and binge eating disorder. Appetite. 2006;46(1):26–30.

43. Zeltser LM. Feeding circuit development and early-life influences on future feeding behaviour. Nat Rev Neurosci. 2018;19(5):302–316.

44. Kim J, Zhang X, Muralidhar S, LeBlanc SA, Tonegawa S. Basolateral to Central Amygdala Neural Circuits for Appetitive Behaviors. Neuron. 2017;93(6):1464-1479.e1465.

45. Sah P, Lopez De Armentia M. Excitatory synaptic transmission in the lateral and central amygdala. Ann N Y Acad Sci. 2003;985:67–77.

46. Beier KT, Steinberg EE, DeLoach KE, et al. Circuit Architecture of VTA Dopamine Neurons Revealed by Systematic Input-Output Mapping. Cell. 2015;162(3):622–634.

47. Jennings JH, Rizzi G, Stamatakis AM, Ung RL, Stuber GD. The inhibitory circuit architecture of the lateral hypothalamus orchestrates feeding. Science. 2013;341(6153):1517–1521.

48. Cowley MA, Smith RG, Diano S, et al. The distribution and mechanism of action of ghrelin in the CNS demonstrates a novel hypothalamic circuit regulating energy homeostasis. Neuron. 2003;37(4):649–661.

49. van den Top M, Lee K, Whyment AD, Blanks AM, Spanswick D. Orexigen-sensitive NPY/AgRP pacemaker neurons in the hypothalamic arcuate nucleus. Nat Neurosci. 2004;7(5):493–494.

50. Aponte Y, Atasoy D, Sternson SM. AGRP neurons are sufficient to orchestrate feeding behavior rapidly and without training. Nat Neurosci. 2011;14(3):351–355.

51. Betley JN, Cao ZF, Ritola KD, Sternson SM. Parallel, redundant circuit organization for homeostatic control of feeding behavior. Cell. 2013;155(6):1337–1350.

52. Yang P, Baker KA, Hagg T. The ADAMs family: coordinators of nervous system development, plasticity and repair. Prog Neurobiol. 2006;79(2):73–94.

53. Blobel CP. ADAMs: key components in EGFR signalling and development. Nat Rev Mol Cell Biol. 2005;6(1):32–43.

54. Weber S, Saftig P. Ectodomain shedding and ADAMs in development. Development. 2012;139(20):3693–3709.

55. Black RA, Rauch CT, Kozlosky CJ, et al. A metalloproteinase disintegrin that releases tumour-necrosis factor-alpha from cells. Nature. 1997;385(6618):729–733.

56. Maretzky T, Schulte M, Ludwig A, et al. L1 is sequentially processed by two differently activated metalloproteases and presenilin/gamma-secretase and regulates neural cell adhesion, cell migration, and neurite outgrowth. Mol Cell Biol. 2005;25(20):9040–9053.

57. Kalus I, Bormann U, Mzoughi M, Schachner M, Kleene R. Proteolytic cleavage of the neural cell adhesion molecule by ADAM17/TACE is involved in neurite outgrowth. J Neurochem. 2006;98(1):78–88.

58. Allinson TM, Parkin ET, Turner AJ, Hooper NM. ADAMs family members as amyloid precursor protein alpha-secretases. J Neurosci Res. 2003;74(3):342–352.

59. Young-Pearse TL, Bai J, Chang R, Zheng JB, LoTurco JJ, Selkoe DJ. A critical function for beta-amyloid precursor protein in neuronal migration revealed by in utero RNA interference. J Neurosci. 2007;27(52):14459–14469.

60. Löffler J, Huber G. Beta-amyloid precursor protein isoforms in various rat brain regions and during brain development. J Neurochem. 1992;59(4):1316–1324.

61. Saftig P, Reiss K. The “A Disintegrin And Metalloproteases” ADAM10 and ADAM17: novel drug targets with therapeutic potential? Eur J Cell Biol. 2011;90(6-7):527–535.

62. Postina R. Activation of α-secretase cleavage. J Neurochem. 2012;120 Suppl 1:46–54.

63. Oyagi A, Moriguchi S, Nitta A, et al. Heparin-binding EGF-like growth factor is required for synaptic plasticity and memory formation. Brain Res. 2011;1419:97–104.

64. Aguirre A, Rubio ME, Gallo V. Notch and EGFR pathway interaction regulates neural stem cell number and self-renewal. Nature. 2010;467(7313):323–327.

65. Idriss HT, Naismith JH. TNF alpha and the TNF receptor superfamily: structure-function relationship(s). Microsc Res Tech. 2000;50(3):184–195.

66. Lawson LJ, Perry VH, Dri P, Gordon S. Heterogeneity in the distribution and morphology of microglia in the normal adult mouse brain. Neuroscience. 1990;39(1):151–170.

67. Ransohoff RM, Cardona AE. The myeloid cells of the central nervous system parenchyma. Nature. 2010;468(7321):253–262.

68. Davalos D, Grutzendler J, Yang G, et al. ATP mediates rapid microglial response to local brain injury in vivo. Nat Neurosci. 2005;8(6):752–758.

69. Nimmerjahn A, Kirchhoff F, Helmchen F. Resting microglial cells are highly dynamic surveillants of brain parenchyma in vivo. Science. 2005;308(5726):1314–1318.

70. Lehnardt S. Innate immunity and neuroinflammation in the CNS: the role of microglia in Toll-like receptor-mediated neuronal injury. Glia. 2010;58(3):253–263.

71. Bessis A, Béchade C, Bernard D, Roumier A. Microglial control of neuronal death and synaptic properties. Glia. 2007;55(3):233–238.

72. Hanisch UK, Kettenmann H. Microglia: active sensor and versatile effector cells in the normal and pathologic brain. Nat Neurosci. 2007;10(11):1387–1394.

73. Minghetti L, Levi G. Microglia as effector cells in brain damage and repair: focus on prostanoids and nitric oxide. Prog Neurobiol. 1998;54(1):99–125.

74. Reemst K, Noctor SC, Lucassen PJ, Hol EM. The Indispensable Roles of Microglia and Astrocytes during Brain Development. Front Hum Neurosci. 2016;10:566.

75. Akiyoshi R, Wake H, Kato D, et al. Microglia Enhance Synapse Activity to Promote Local Network Synchronization. eNeuro. 2018;5(5).

76. Semple BD, Blomgren K, Gimlin K, Ferriero DM, Noble-Haeusslein LJ. Brain development in rodents and humans: Identifying benchmarks of maturation and vulnerability to injury across species. Prog Neurobiol. 2013;106-107:1–16.

