## Supplemental Data for "Prefrontal cortical protease TACE/ADAM17 is involved in neuroinflammation and stress-related eating alterations"

**SUPPLEMENTARY MATERIALS**

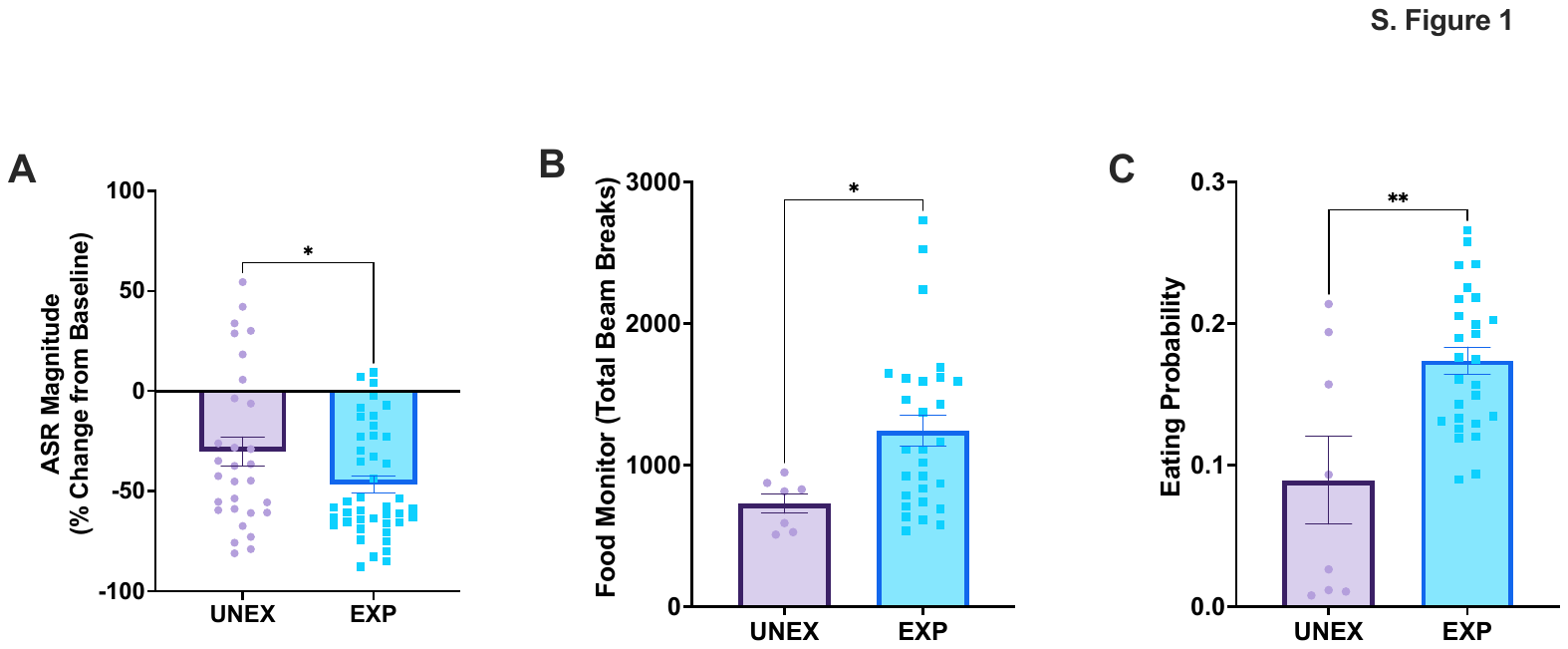

**Supplemental Figure 1. Behavioral alterations in a rat model of adolescent PTSD. (A)** Exposure to predatory stress and isolation during adolescence resulted in blunted acoustic startle reflex (ASR) reactivity (*t*_71_=2.13, *p*=0.037). PTSD, n = 30; unexposed controls, n = 43. **(B-C)** Eating-related behaviors in surgically naïve rats exposed to stress. **(B)** Traumatic stress during adolescence increased the number of IR beam breaks in the food monitor relative to unexposed controls (*t*_33_=2.28, *p*=0.029). **(C)** Trauma increased the probability that rats exhibited eating behaviors relative to controls (*t*_33_=3.50, *p*=0.0014). Surgically naïve rats: unexposed controls, n = 8; PTSD, n = 27. *, *p*<0.05; **, *p*<0.01.

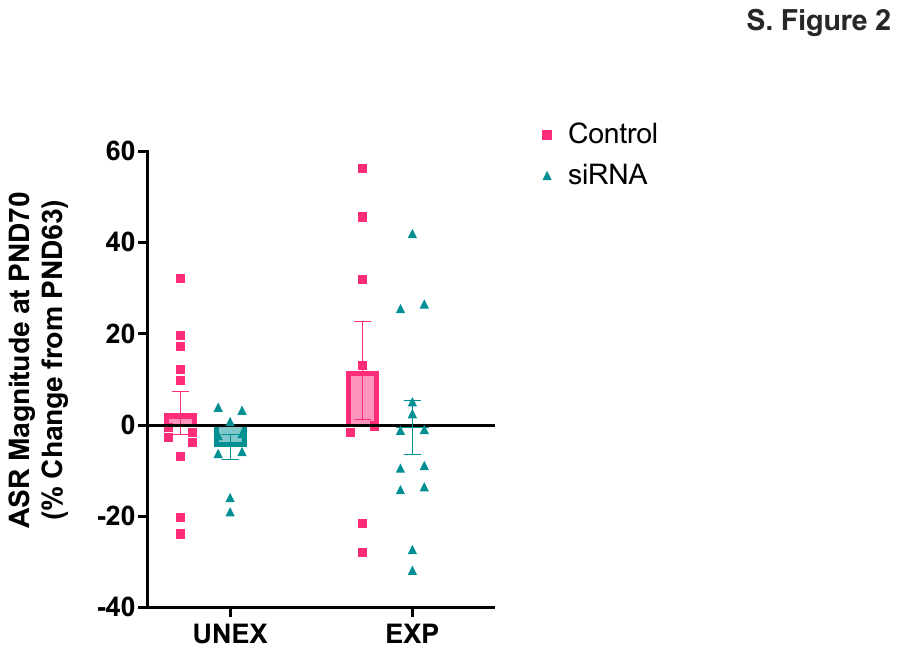

**Supplemental Figure 2. Effects of intracerebral TACE/ADAM17 siRNA injections on the acoustic startle.** ASR responses were unaltered at one-week post-siRNA administration. The bar graph illustrates the ASR magnitude at PND70 as the percent change from PND63 (post-stress). Stress: F_1, 38_=1.17, *p*=0.29; Treatment: F_1, 38_=2.42, *p*=0.128; Interaction: F_1, 38_=1.55, *p*=0.70. Control Unexposed, n = 12; siRNA Unexposed, n = 9; Control Exposed, n = 8; siRNA Exposed, n = 13.

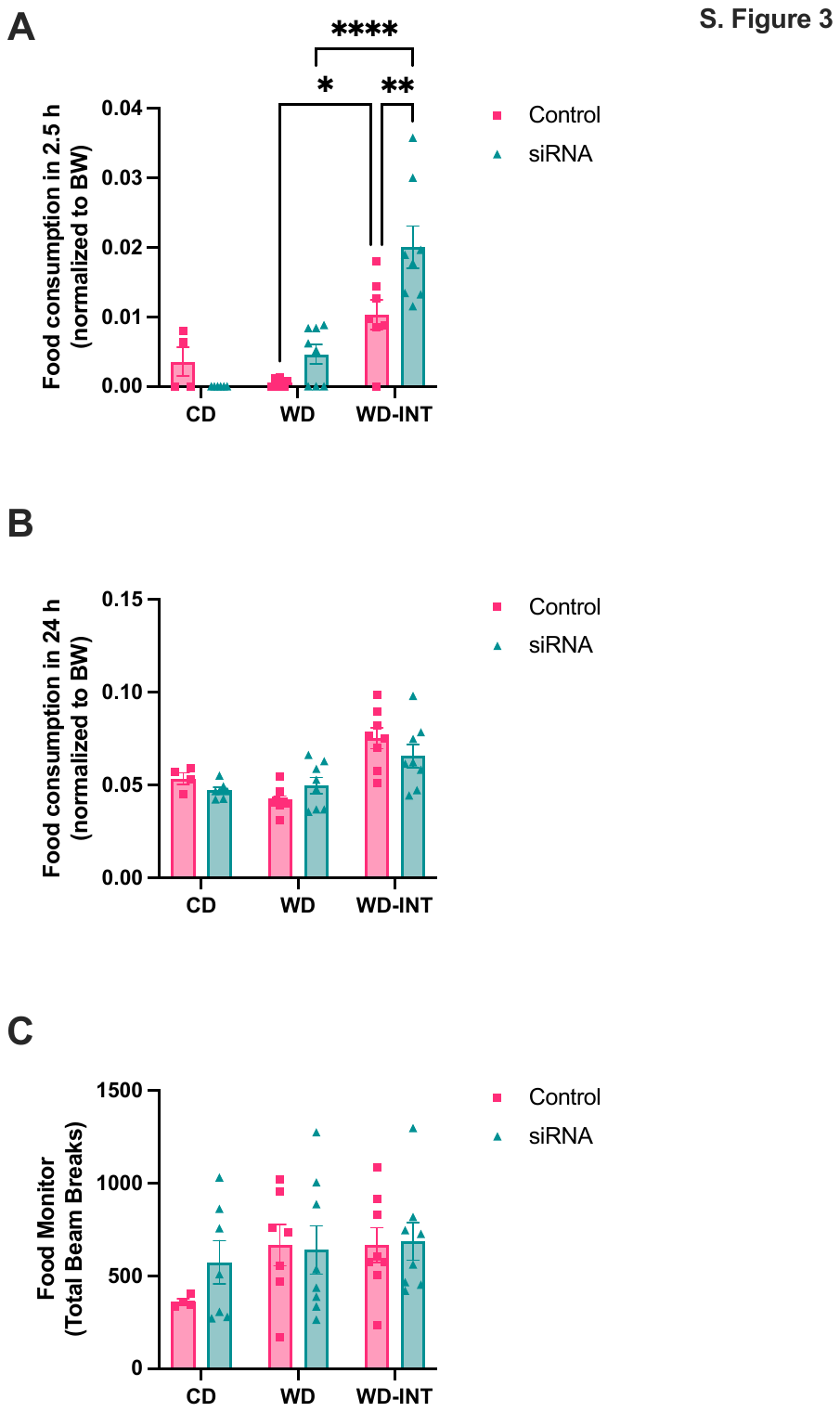

**Supplemental Figure 3. Food intake alterations in rats exposed to an intermittent obesogenic high-fat diet.** A subgroup of rats was introduced to a Western-like obesogenic diet for 48 h (same obesogenic diet used in our prior studies, Vega-Torres et al., 2018, 2022). The rats were re-introduced to the diet once a week for three consecutive weeks to generate a well-established model of binge eating (Czyzyk et al., 2010; Maske et al., 2020). **(A)** Weight-normalized food consumed during 2.5 h following the third experimental cycle. As expected, WD-INT rats consumed more food than CD and WD rats (Diet: F_2, 34_=29.96, *p*<0.0001; Treatment: F_1, 34_=4.27, *p*=0.046; Interaction: F_2, 34_=5.01, *p*=0.012). TACE/ADAM17 siRNA administration increased binge eating-like behavior relative to vehicle controls (*p*=0.0091, difference: -0.0097, 95% CI of difference: -0.018 to -0.0018). **(B)** Weight-normalized food consumed during the 24-h period in cycle 3. Diet: F_2, 37_=17.74, *p*<0.0001; Treatment: F_1, 37_=0.47, *p*=0.50; Interaction: F_2, 37_=2.25, *p*=0.12). Twenty-four h food intake was similar between WD-INT siRNA and control groups, confirming the binge eating phenotype at 2.5 h (*p*=0.63). **(C)** The total number of nose pokes inside the food monitor was similar between groups. Diet: F_2, 36_=1.74, *p*=0.19; Treatment: F_1, 36_=0.53, *p*=0.50; Interaction: F_2, 36_=0.52, *p*=0.60). CD, control diet; WD, Western-like high-fat diet, WD-INT, WD intermittent access. WD-INT control injection, n = 8; WD-INT siRNA injection, n = 8. *, *p*<0.05; **, *p*<0.01; ****, *p*<0.0001.
